# Noise Propagation in Transcriptional Genetic Cascades

**DOI:** 10.1101/2023.09.17.558128

**Authors:** JD Marmolejo, JM Pedraza

## Abstract

Cellular processes are inherently stochastic, leading to fluctuations in protein concentration quantifiable as noise in gene expression. Precise description of noise propagation in gene networks is essential for designing noise-tolerant gene circuits and understanding signal reliability in biological networks, but current models for noise propagation are primarily system-specific or limited to short gene cascades. Here we present an analytical expression for noise propagation in gene expression that works for long cascades and incorporates global noise. Since modelling all aspect of noise can be prohibitively complicated, in many situations only intrinsic or global noise is considered, but general criteria for when each type is dominant are still lacking. As an example of the possible use of our analytical expression, we examine the role different aspects of the network have on the balance between intrinsic and global noises and their propagation. We show that the type of cascade, cascade length, sensitivity, and basal transcription rates have an effect beyond simple protein abundance. This has practical implications for designing synthetic gene networks in prokaryotes and improving our knowledge on noise propagation in gene networks, and could shed light on how evolution may shape circuit sizes to balance signal fidelity and metabolic cost.

## 1 Introduction

Information processing within cells relies on the complex network of chemical reactions that constitutes gene expression. Proper functioning of these processes can be limited by signal fluctuations throughout these networks. These signal fluctuations are inevitable because gene expression is inherently a stochastic process [1–4]. Stochasticity in gene expression thus has a dual impact, potentially hindering intracellular signaling due to protein level fluctuations, while also generating phenotypic diversity that enables bacterial adaptation and survival [5]. In transcriptional gene circuits, signal variations due to stochasticity are called simply “noise in gene expression”, and are measured as the differences in mRNA and protein quantity between identical cells [3]. It is usually quantified as the coefficient of variation (CV). It can be classified functionally as intrinsic or extrinsic [1] when for two copies of the same gene their fluctuations are uncorrelated (intrinsic) or correlated (extrinsic), or as intrinsic, transmitted and global [6], depending on whether its origin lies in the individual events of transcription, translation and degradation (intrinsic), in the fluctuations of other genes in the network (transmitted) or of other cellular components involved in gene expression such as fluctuations in polymerases, ribosomes or cell growth (global). Transcriptional networks are usually examined using stochastic simulations or analytically by system-specific models. General models to describe noise propagation have been developed using a master equation approach [7], but without global noise because its origin has not been elucidated to the point where you can incorporate it into such models. A Langevin equation approach [8] has been used which allows for a phenomenological incorporation of global noise but only for short cascades [6]. Here we present a mathematical generalization for the model of noise propagation in transcriptional networks, which is useful for predicting fluctuations in transcriptional cascades of any length, and whether they are composed of repressors, activators, or both. Then we show via numerical assessments of our model that, particularly in repressor cascades, the magnitude of total noise and the equilibrium between extrinsic and intrinsic noises are especially impacted by the sensitivity to repression of each gene in the network, its basal transcription rate and its length. The noise behaviour as a function of the circuit length offers a valuable insight into the reason why certain transcriptional cascades tend to be short and why transcriptional circuits are modularized into smaller parts. When modeling noise in experimental contexts, it can be practical to use only global or intrinsic factor depending on whether the measurements indicate only one dominates, but there is no general rule of thumb for knowing when the gene in a specific circuit is mostly subject to global or intrinsic noise. Our approximate but analytical results offer a quick way of estimating their relative importance based on the biochemical parameters, which can then be used to justify simpler models. Taking into account the stochastic properties of gene networks, both in the design of synthetic circuits and in the analysis of natural ones, is desirable but will only happen if the models required do not become unmanageably complex mathematically or prohibitively expensive computationally. Our results should provide a tool for guiding the construction of case-specific, approximate models that are useful but practical.

## 2 Theoretical results

Following [6, 8–10], we used the Langevin approach because it allows for sequential solution for a cascade gene network and because it allows for the incorporation of an effective noise term. In contrast to the model in [6], in our model the global fluctuations have an autocorrelation that decays exponentially [11, 12], since protein-based components of the expression machinery fluctuate with a correlation dependent on their degradation/dilution rate [1] and size and growth dynamics can affect noise in gene expression [13]. Here we will show the main results of our study; for more details see the Supplementary Information and our GitHub repository.

### 2.1 Initial considerations

We considered only cascades, genetic circuits in which each gene exerts control solely over the gene immediately downstream from it. We define *δx* = *x x*, the variation of x with respect to its expected value, where x is a random variable. Using separation of timescales, we eliminate the equations for the mRNA dynamics, to obtain the following equation for the proteins expressed from the *i-th* gene in the cascade

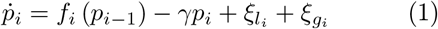

where *i >* 1, *f*_*i*_ (*p*_*i*−1_) is a Hill function, *γ* is the protein degradation rate, 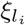 corresponds to the local or intrinsic fluctuations and 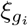 represents the fluctuations due to global variations in the concentration of cellular components. We assumed *f*_*i*_ (*p*_*i*−1_) is proportional to a Hill function, shown in equation (2), since it is the preferred model for these systems; however, our model is general enough to consider other possible functions for the interactions. We considered that all protein degradation rates are approximately equal to the cell growth rate, because in stable proteins, the protein degradation rates are mostly determined by the rate of dilution that occurs as a result of cell division [14].

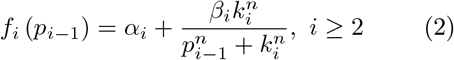

Regarding the statistical properties of 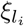 and 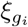, we considered that 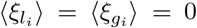, that 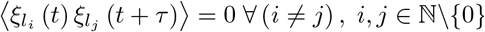, *i, j* ∈ N\{0}, and that 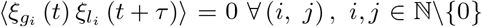, *i, j* ∈ N\{0}. We defined

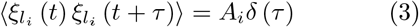

where *A*_*i*_ ∈R^+^. This captures the independence of transcription and destruction events, which is a good approximation given that the degradation rate for proteins is lower than the degradation rate for the mRNA and that many proteins are produced per mRNA [6]. It is worth noting that the factor *A*_*i*_ now depends on the burst size (average number of proteins produced by an mRNA during its lifetime).

We also defined

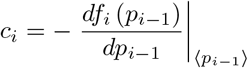

where *i* ≥ 2, which can be written as

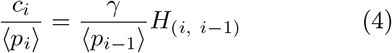

where

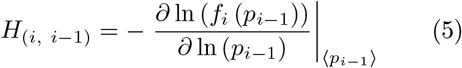

is the logarithmic gain of gene *i*, given the control exerted by gene *i-1*. If the *i-th* gene is controlled by a repressor, *c*_*i*_ *>* 0 because the Hill function will be a monotonically decreasing function. For the case of an activator *c*_*i*_ will have the opposite sign. The equation (4) will be valid initially for *n* ≥ 2 and by definition *c*_1_ = 1. The fact

### 2.2 Model

We assume that global fluctuations will affect the expression rates multiplicatively [6] and that the global noise will have an autocorrelation given by that we will work only with this first order derivative means we are using a linear approximation around the steady state of each gene, and is the strongest approximation in our model.

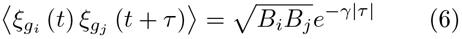

where 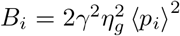 and 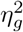 represents the global noise magnitude, a parameter inferred from experiments. From equation (1), if we evaluate in the *n-th* gene, compute the variation of *p*_*n*_ around the steady state —which coincides with the expected value—, and perform the Fourier Transform, we obtain that

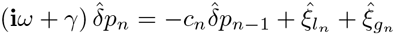

where 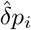 denotes the Fourier Transform of *δp*_*i*_. It is possible to prove by induction (see Supplementary Information) that this expression is equivalent to equation (7).

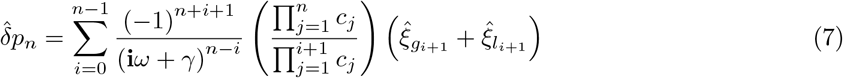

Defining

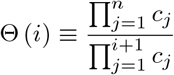

multiplying by the complex conjugate and averaging we obtain,

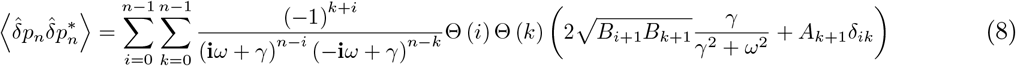

Also defining,

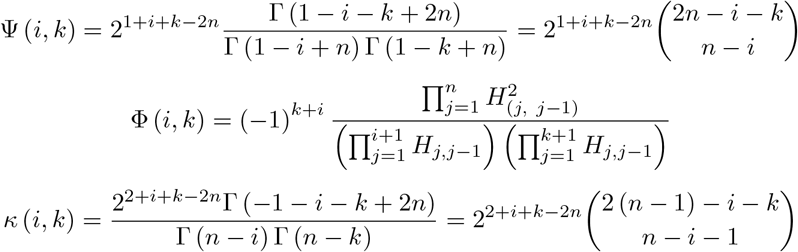

using the Wiener-Khinchin theorem, and dividing by ⟨*p*_*n*_ ⟩^2^, we obtain that

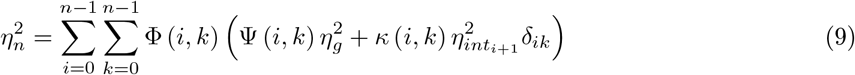

The noise in gene expression of the *n-th* gene of the cascade.

This approximation also implies the appearance of coefficients associated with time averaging, as reported in [7], but as shown in equation (9), where they take the form of binomial coefficients. On the other hand, because the calculations were made linearly around the steady state, the model will only be valid within the bulk of the distribution [6]. As we said before, our results are not limited to cascades composed solely of activators or repressors, because the nature of the transcriptional control exerted by the genes will be embedded in the sign of the logarithmic gain, which allows our model to be valid for every cascade configuration as long as there are not behaviors such as feedback loops. The model could in principle be extended to such cases, but the analytical calculations become much more difficult and limited because the possible existence of multiple steady states reduces the validity of the local linear approximation.

When substituting into equation (9), if we replace *n* with 1, 2 and 3 we have

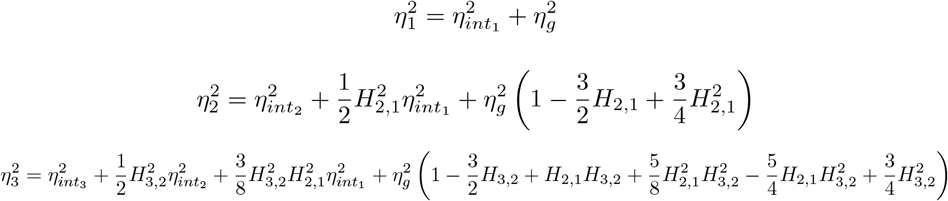

Which coincides with the results of [6] except for the numerical coefficients which now correspond to the time averaging of colored noise instead of white noise for the global fluctuations.

## 3 Numerical evaluation of model parameters

In order to use our analytical expression to explore the behavior of both types of noise in generic cascades, we explored parameter space by assigning random parameters 3.0 × 10^4^ times, grouped by Hill coefficient ranges, cascade lengths and basal transcription rates. To ensure statistical validity and sufficient sampling, the selection was limited to cascades with at most 6 genes, as longer cascades would require significant additional computational resources and time. Although this approach has a trade-off between feasibility and accuracy, it is important to note that a 6-gene cascade is still relatively large compared to typical systems studied in the literature.

In general, uniform distributions were fixed between 1*/*8 *min*^−1^ and 1*/*3 *min*^−1^ for mRNA degradation rate [15]; between 1*/*60 *min*^−1^ and 1*/*18 *min*^−1^ for protein degradation rate, since bacterial doubling times in rich media are usually in the range [18*min*, 60*min*] depending on the culture medium, temperature, species, and other factors; between 10^3^ and 10^4^ —the typical orders of magnitude for amount of proteins—, for *k*_*i*_, given that we are considering Hill functions for our numerical evaluations. The magnitude *η*_*g*_ was estimated at 0.25. Given that the closer the Hill coefficient is to 0, the more the genes will behave as constitutive genes, the minimum value for it was fixed at 0.5.

From now on we will call 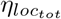 or local noise, the intrinsic noise in the last gene of the cascade plus the transmitted effect of the intrinsic noises of upstream genes. Similarly, 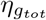 will represent the effect of global fluctuations, including both the direct effect on the last gene and the transmitted effect from the previous genes. Bearing in mind that basal transcription rate can affect the control level of upstream genes over those downstream, we have evaluated basal transcription rates (*α*) equal to 0.1*β*, 0.15*β* and 0.2*β*, where *β* is the maximum transcription rate.

Equation (9) shows that noise propagation depends sensitively on the Hill coefficient *h*, present within the logarithmic gains. Since we want to evaluate its importance in the balance between intrinsic and global noise, we want to explore multiple values for the different genes on the cascade, but to separate the effect of high values of *h* from high variability in *h* within a cascade, we have performed evaluations with two types of uniform probability distributions for *h*. We have examined the case where *h* ∼ U (0.5 + *i*, 0.6 + *i*), *i* ∈ {0, 0.1, 0.2, …, 2.9}, where 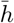 corresponds to the middle point of each [0.5 + *i*, 0.6 + *i*] interval, and where *h* ∼*u* (0.5, *i*), *i* ∈{0.6, 0.7, …, 3.5}, Δ*h* = *i* − 0.5. We compared the noise behavior of gene cascades with different magnitudes and levels of diversity in sensitivity to upstream gene control, and check whether the observed phenomena are consistent across these scenarios. Our results are shown in Figures 1 and 2 and both graph-ics show 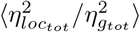 in the ordinate axis. Figures 3 and 4 show, in contrast, average total noise 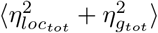

**Figure 1:**
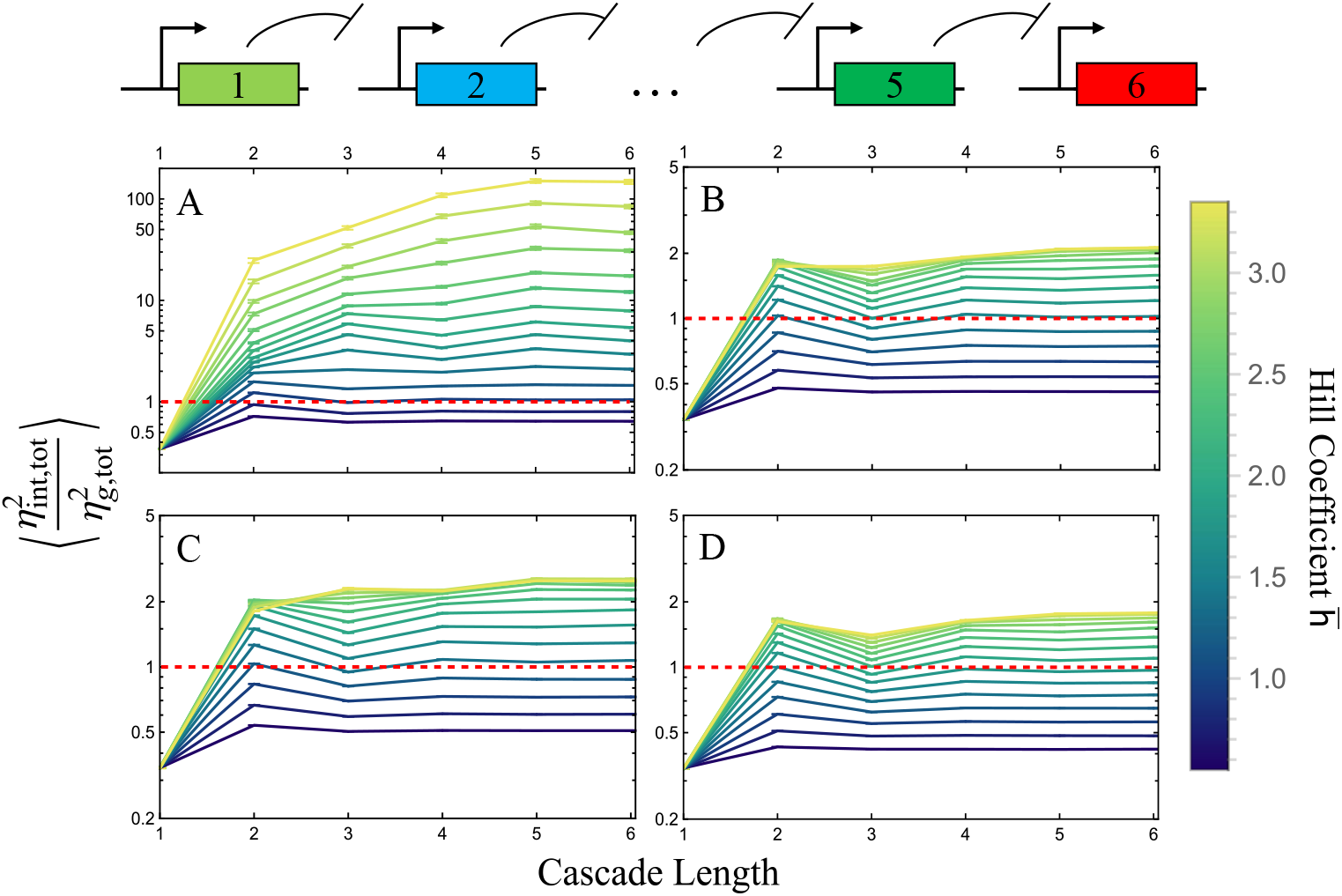
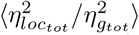 as a function of the cascade lengths for repressor-only cascades. Here we considered *h* ∼*u* (0.5 + *i*, 0.6 + *i*), *i* ∈ {0, 0.1, 0.2, …, 2.9}. Each point of the plot is the average of 3.0 × 10^4^ generated data. The lines connecting the points are included for the purpose of facilitating visualisation. Error bars show the standard error of the mean. The fixed red dashed line shows where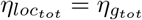. All plots are presented in logarithmic scale. The top scheme depicts the network in which we primarily operate. Figure (A) shows systems with *α* = 0, (B) considering *α* = 0.1*β*, (C) *α* = 0.15*β* and (D) *α* = 0.2*β*. Note the different scale on the y-axis in Figure (A). Oscillation-like behavior is related to gene positions, since even genes are repressed and odd lengths are expressed.

**Figure 2:**
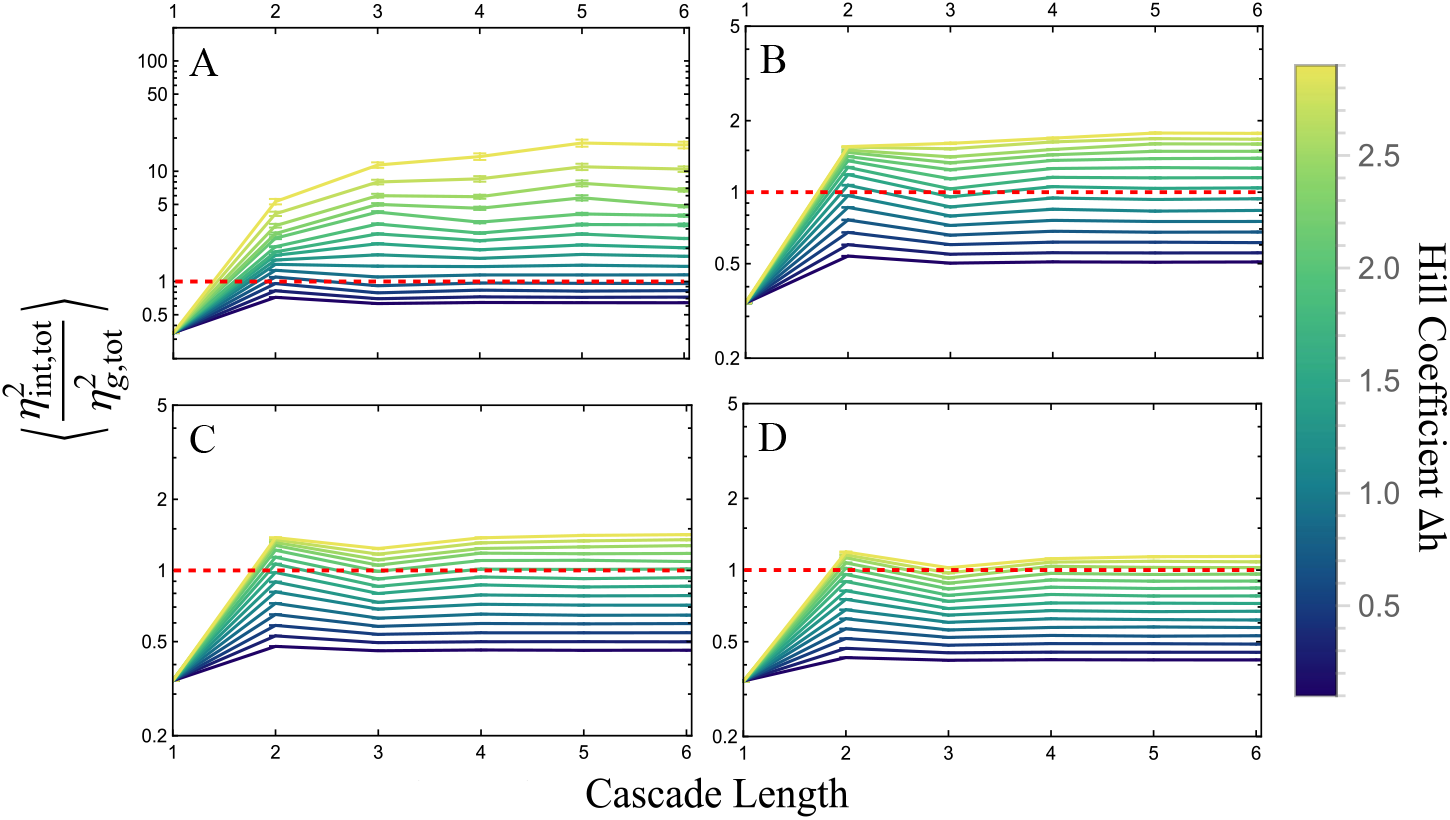
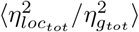as a function of the cascade lengths for a moving window of *h*. We considered*h* ∼ *u* (0.5, *i*), *i* ∈ { 0.6, 0.7, …, 3.5}, Δ*h* = *i* − 0.5, so that *h* ∈ [0.5, *i*]. Each point of the plot is the average of 3.0 × 10^4^ generated data and error bars show the standard error of the mean. The lines connecting the points are included for the purpose of facilitating visualisation. The fixed red dashed line shows where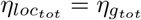. All plots are presented in logarithmic scale. As in Figure 1, (A) achieves higher values and has a different scale. Figure (A) shows systems with *α* = 0, (B) with *α* = 0.1*β*, (C) *α* = 0.15*β* and (D) *α* = 0.2*β*.

Here we discuss the numerical evaluations of our models for repressor-only cascades; however, simulations of systems including activators were carried out in order to evaluate the behavior of the noises in such systems as well (see Supplementary Information).

## 4 Discussion

The usefulness of an analytical expression for the noise in a gene network can be limited by the choice of biological details to ignore, the mathematical approximations used, the imprecision in the biochemical parameter measurements, and in our model the fact that the global noise parameter is inferred from measurements in a way that depends on the model itself. However, having an analytical expression at hand allows for the rapid exploration of parameter space. In particular, we are interested not only on the total noise but on the balance between intrinsic and global noise for different configurations of the cascade. When the classification of noise types into intrinsic and global noise began, it was widely believed that global fluctuations were often the main source of noise in gene expression, at least for common conditions for *E. coli* [16] and *S. cerevisiae* [17]. In addition, some studies indicated that the balance between intrinsic and global noise depended mostly on protein abundance, as proposed in [16]. In [18], the authors suggested that for intermediate protein abundances in *S. cerevisiae*, the noise in gene expression is not completely dominated by global factors, but is comparable or even surpassed by gene-specific noises. Moreover, according to [14], in *E. coli* intrinsic noise is dominant for low protein concentrations and global noise is dominant in the opposite case. On the other hand, [8] used analytical results (on which our method is based) to examine the effect of cascade length on the propagation of intrinsic noise. Our approach allowed us to test these ideas for multiple parameters, including global noise, and observe behaviors similar to those previously reported, as well as evaluating the relative importance of the noise sources as a function of the biochemical parameters such as the sensitivities and basal transcription rates.

### 4.1 Basal transcription rates may affect the noise balance and propagation

Transcription at a low basal rate may occur even when a gene is repressed by a fully bound transcription factor, and even when there is a high concentration of repressor the stochasticity of binding and unbinding means that a fraction of the time the promoter might be unbound. These factors result on a basal transcription rate. Our results suggest that the relative ratio of basal to maximum transcription rate plays a critical role in regulating the balance between intrinsic and global noise in repressor cascades.

According to our analysis, it is possible for intrinsic noise to overcome global noise in repressor-only cascades; however, when the basal transcription rate *α* increases relative to the maximum transcription rate *β*, the possibility of intrinsic noise exceeding global noise decreases. This is in line with literature, as an increase in *α* relative to *β* results in genes becoming less sensitive, even for high *h*, leading to higher protein levels and a more dominant global noise. This means that sacrificing sensitivity for stability via the basal rate is a limited strategy because the reduction in total noise is bound by the global noise. However, as seen in Figure 4, there is an important reduction in total noise when initially increasing the basal transcription rate from 0.

### 4.2 The significance of protein abundance for noise balance decreases with cascade length and gene sensitivity

Another factor that has been postulated to determine the balance between types of noise is the protein abundance, as stated in [14] for *E. coli* and [18] for *S. cerevisiae*. It has been established in early studies [19] that noise decreases inversely with protein amount. Nevertheless, our results show that global noise is dominant in constitutive genes and insensitive circuits, but intrinsic noise can increase its dominance as sensitivity increases, regardless of protein abundance. Furthermore, the balance between intrinsic and global noise becomes increasingly independent of protein amount as network size increases, both for very low and very high sensitivity levels.

Figures 1 and 2 show that, in general, small sensitivities result in global noise being dominant, regardless of cascade length and basal transcription rates. In contrast, as sensitivity increases, there is a shift in the balance of intrinsic and global noise, with intrinsic noise becoming increasingly dominant. This effect could be a consequence of an increase in the logarithmic gain value, which results in a greater importance of genes further upstream. This effect is key in ensuring the dominance of intrinsic noise over global noise, as intrinsic noise increases more than global noise as upstream genes become increasingly important.

On the other hand, the cascade length parity (odd or even numbers) appears to have diminishing effects in noise balance as the length and sensitivity increase, particularly because oscillatory behavior is less observable and because intrinsic noise tends to be dominant over global noise. Since the parity is directly related to protein abundance, noise balance could be less dependent on protein quantity as sensibility increase in the network.

The attenuation of oscillatory-like behavior in the noise ratio is more pronounced as the length of the network increases, regardless of parity, because noise ratios tend to stabilise with increasing cascade length as shown in Figures 1 and 2, particularly for very low and very high sensitivities. One hypothesis to explain this effect is that, at low sensitivities, the spread of noise from upstream genes further away is less important, and the balance of noise is predominantly based on nearest upstream neighbours, which could lead to a constant ratio. For higher sensitivities, longer cascades tend to have a noise ratio that is intermediate between 2and 3-cascades depending on the sensitivity level, which could be attributed to the increasing importance of upstream genes. As sensitivities increase, an equilibrium-like behavior is achieved, and oscillations are lost once again, possibly due to a limit on the propagation of both types of noise.

Additionally, other systems like activators-only and mixed cascades (see Supplementary Information) may have different behaviors. Our simulations showed that the evaluated systems of activators-only cascades tend to have a global noise dominance. This may be an effect of the protein abundance, since every gene in an activator cascade tends to be highly expressed.

### 4.3 Noise balance could affect evolution of cascade lengths

The results of [8] indicated that amplification cascades were not limited by noise in the sense that high amplification and reasonable stability could be obtained even for long cascades. We are interested in the more general case of distinguishing among many possible states, and our evaluation of widely varying sets of parameters indicates that, except for low sensitivities or high basal transcription rates, the total noise does on average increase with cascade length.

Results depicted in Figures 3 and 4 indicate that shorter repressor-only cascades, on average, exhibit lower total noise than longer cascades, whose total noise increases significantly as sensitivity on the network increases. This effect is particularly pronounced in the case of networks with low basal transcription rates. This finding is in line with what we have previously described in section 4.1. Additionally, insensitive repressor cascades tend to be noisier than sensitive ones when considering diverse *h* and high basal transcription rates, probably because sensitive networks are more affected by length parity.

**Figure 3:**
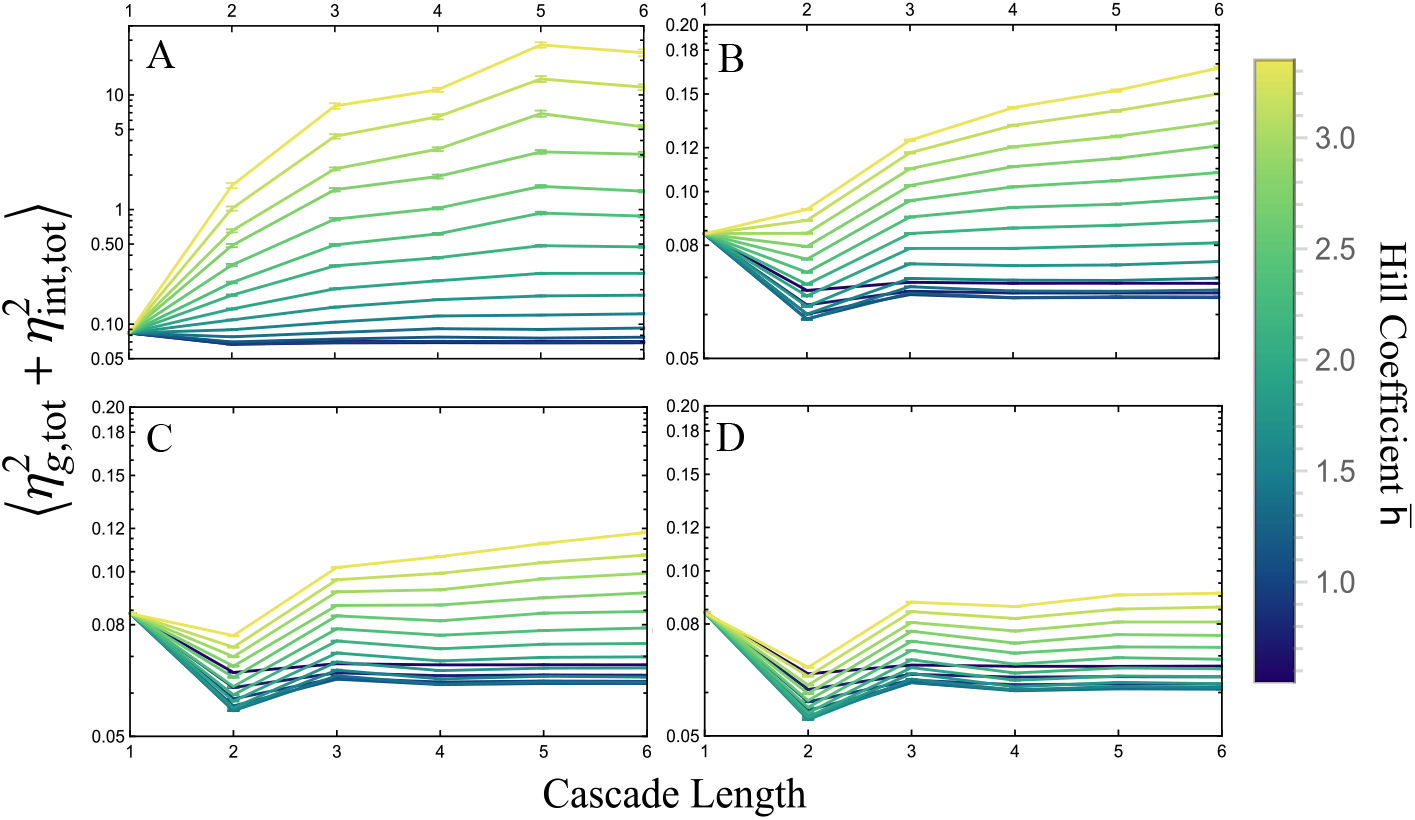
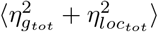 as a function of the cascade lengths for a varying width of *h* values. We have employed the same probability distribution as depicted in 1. Each point of the plot is the average of 3.0 × 10^4^ generated data and error bars show the standard error of the mean. The lines connecting the points are included for the purpose of facilitating visualisation. All plots are presented in logarithmic scale. As in Figure 1, (A) achieves higher values and has a different scale. Figure (A) shows systems with *α* = 0, (B) with *α* = 0.1*β*, (C) *α* = 0.15*β* and (D) *α* = 0.2*β*.

**Figure 4:**
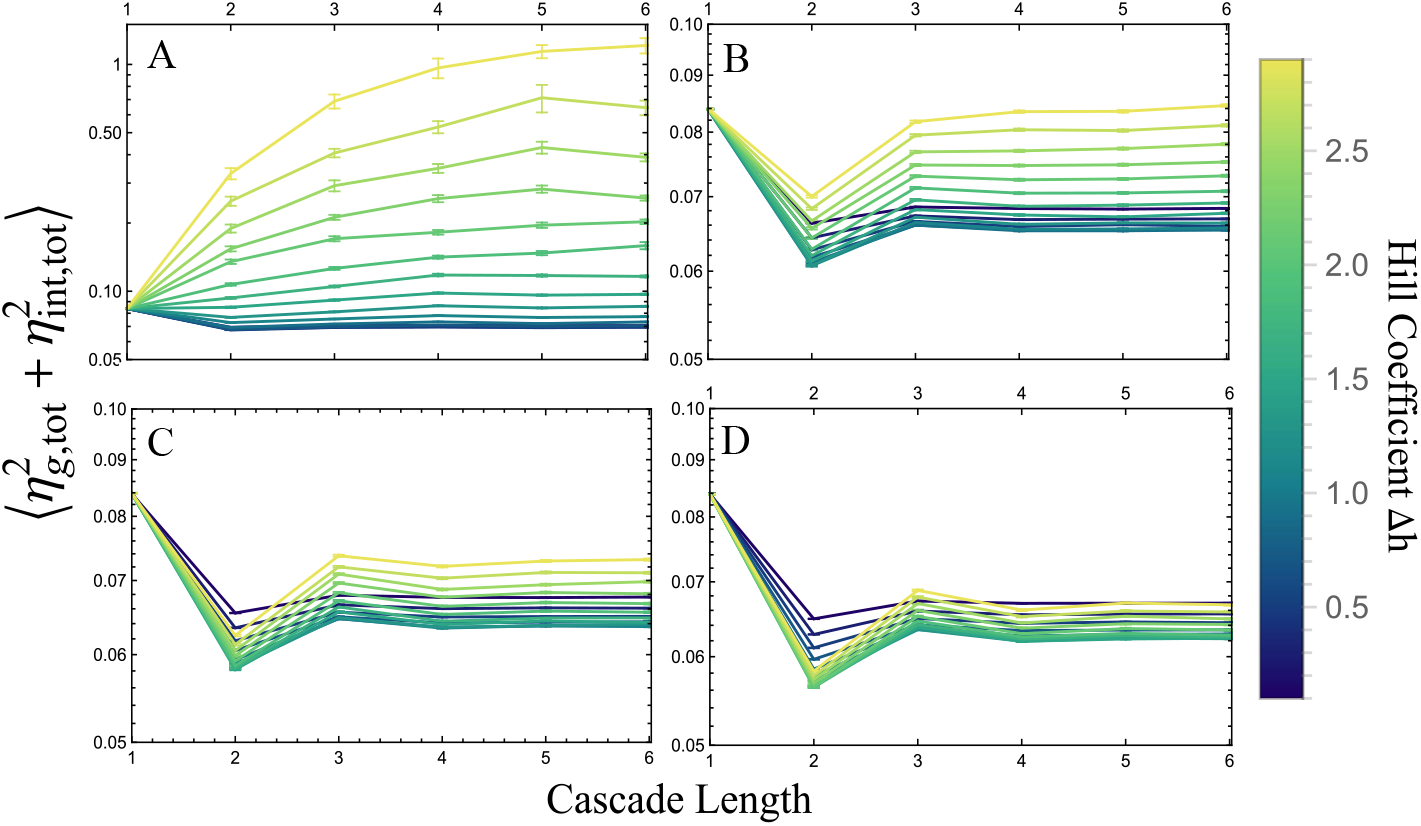
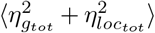 in function of the cascade lengths for repressor-only cascades. We have employed the same probability distribution as depicted in 2. Each point of the plot is the average of 3.0 × 10^4^ generated data and error bars show the standard error of the mean. The lines connecting the points are included for the purpose of facilitating visualisation. All plots are presented in logarithmic scale. Figure (A) achieve higher values than the others and scales are different than those shown in Figure 3 for visualisation purposes. Figure (A) shows systems with *α* = 0, (B) with *α* = 0.1*β*, (C) *α* = 0.15*β* and (D) *α* = 0.2*β*.

Our numerical evaluations of equation (9) show that, on average, shorter repressor-only cascades may be more useful for the cell when responsiveness and signal reliability are important, just as it happens in the detection or response to continuously varying environmental stimuli, since total noise tend to be lower in high sensitivity conditions. Long cascades may be more useful for the cell for performing computations with lower need of signal reliability or digital-like outputs. However, metabolic load considerations can also play a role in their evolution, as a shorter cascade of highly expressed genes can be more costly than a longer cascade of mildly expressed ones.

## 5 Conclusions

We presented a generalized model, based on the analytical methods of [8] and the inclusion of global noise presented in [6], which allows us to predict the noise in gene expression for an arbitrary cascade composed by *n* genes. In contrast with [6], we consider an exponentially decaying autocorrelation for global fluctuations. Our result, while approximate, can be used to quantify noise in genetic cascades utilising a compact formula, which is useful to avoid simulations and save calculation time. It also provides the possibility of evaluating whether to only consider global or intrinsic noise sources when constructing an approximate model for a specific system without having to perform direct experimental measurements.

As an example of its usefulness, we numerically evaluated our model for multiple combinations of parameters, examining the balance between intrinsic and global noise as factors such as cascade length, sensitivities, basal transcription rates, and the network’s overall architecture. Our assessments indicate that repressor-only cascades have a dominant total global noise for low sensitivity levels, but not for longer and more sensitive cascades, regardless of protein abundance. The impact of protein abundance may thus be overshadowed by the propagation of noise from upstream genes. Furthermore, it is possible that transcriptional networks have evolved to be shorter to achieve higher sensitivity for analog stimuli detection or response, while longer networks may have evolved to perform digital operations that do not require high signal reliability.

Our study highlights the importance of considering the type of noise for effective control of gene expression fluctuations, in both natural prokaryotic systems and synthetic circuit design. The behaviour and propagation of noise in transcriptional cascades depends on the specific network topology and system parameters, but we identified some trends that could guide understanding of these systems. We hope that the availability of an easy to evaluate analytical expression for the noise will help incorporate stochasticity considerations in the design of synthetic gene circuits and the analysis of naturally occurring transcriptional signalling cascades.

## Acknowledgments

We express our gratitude to J MejÍa for providing valuable advice on simplifying the calculations, and to S Henao and the Systems Biology Uniandes group for engaging in fruitful discussions.

